# Haplotype-specific expression of a terpene synthase underlies linalool variation in the grapevine cultivar Riesling

**DOI:** 10.64898/2026.04.26.720724

**Authors:** Jerry Lin, Mélanie Massonnet, Noé Cochetel, Larry Lerno, Luis Diaz Garcia, Mireia Domenech-Lopez, Rosa Figueroa-Balderas, Susan E. Ebeler, Dario Cantu

## Abstract

Grapevine cultivars vary widely in monoterpenoid content, yet the genetic and regulatory mechanisms underlying this variation remain poorly characterized beyond highly aromatic Muscat types. We profiled free volatiles and monoterpenoid glycosides in a Riesling × Cabernet Sauvignon F1 mapping population, revealing extensive variation and transgressive segregation consistent with multigenic control. QTL mapping identified 59 significant loci associated with 47 volatile compounds and monoterpene glycosides, including two major QTLs explaining 33.8% and 32.4% of phenotypic variance in (3*S*)-linalool accumulation. Integration of haplotype-resolved transcriptomics with metabolite data, enabled by a chromosome-scale diploid Riesling genome assembly, resolved a (3*S*)-linalool/nerolidol synthase cluster on chromosome 10 and identified *VviTPS54* as the strongest candidate underlying linalool variation. *VviTPS54* exhibited haplotype-specific expression strongly correlated with (3*S*)-linalool accumulation across genotypes, while no QTL was detected at the *VviDXS1* locus and *VviDXS1* expression was not correlated with terpene levels, indicating that regulatory variation within terpene synthase clusters, rather than methylerythritol phosphate (MEP) pathway flux, drives monoterpenoid composition in this population. These results establish regulatory variation of terpene synthases as a key mechanism underlying monoterpenoid diversity in grapevine and demonstrate that resolving such variation requires haplotype-phased genome assemblies coupled with haplotype-resolved transcriptomics to detect allele-specific expression differences at complex, heterozygous loci.

## Introduction

Disease pressure has driven the introgression of resistance traits from wild grape species into domesticated grape (*Vitis vinifera* ssp. *vinifera*; hereafter *V. v. vinifera*) backgrounds, while ongoing and projected climatic challenges have renewed interest in traits such as drought tolerance, heat tolerance, and altered phenology (Cantu *et al*., 2024). However, wild germplasm often introduces undesirable aroma and flavor attributes, and its effects on the volatile profile and resulting organoleptic properties of fruit and wine remain difficult to predict (Yang *et al*., 2016). For example, crosses of *V. v. vinifera* with *V. x labruscana* yield progeny with elevated ester levels absent in *V. v. vinifera*, while hybrids with *V. amurensis* and *V. thunbergii* show reduced volatile diversity (Yang *et al*., 2009). Different accessions of *V. cinerea* and *V. riparia* exhibit wide variation in methoxypyrazine content, further affecting their utility in breeding programs (Sun *et al*., 2011). Compounding this challenge, grapevine genomes are highly heterozygous and harbor recessive deleterious alleles, which complicate the recovery of desirable traits through backcrossing (Zhou *et al*., 2017). Identifying quantitative trait loci (QTLs) and tightly linked genetic markers for key organoleptic traits would enable early selection of seedlings, accelerate breeding cycles, and improve parental selection strategies.

Volatile aroma compounds are key determinants of wine sensory properties that arise from grapes, microbial metabolism during fermentation, and aging and storage conditions (Robinson *et al*., 2014). Grape-derived aroma compounds can largely be grouped into eight major classes based on biosynthetic origin: lipoxygenase pathway products (e.g., C6 volatiles such as green leaf aldehydes), monoterpenoids, sesquiterpenoids, norisoprenoids, furan derivatives (e.g., furaneol), methoxypyrazines, volatile sulfur compounds, and phenylpropanoid-derived compounds (Robinson *et al*., 2014; Lin *et al*., 2019).

Compared to fruits such as apples, strawberries, and peaches, most grape species have relatively low levels of esters and terpenoids. Notable exceptions include *V. x labruscana*, whose aroma is characterized by esters such as methyl anthranilate, and “aromatic” *V. v. vinifera* cultivars that accumulate higher concentrations of monoterpenoids (Sun *et al*., 2011). Within *V. v. vinifera*, cultivars can be broadly classified into three groups based on monoterpenoid content: “neutral” or “non-aromatic” cultivars (e.g., Chardonnay, Cabernet Sauvignon) with very low levels; intermediate “aromatic” cultivars such as Riesling (1-4 mg/L); and highly aromatic Muscat cultivars, which can exceed 6 mg/L of total monoterpenoid content (Rapp, 1998; Mateo and Jiménez, 2000).

Monoterpenoids are among the most important contributors to grape and wine aroma. Even in cultivars not typically classified as aromatic, monoterpenoids, together with related terpenoids such as sesquiterpenoids and norisoprenoids, contribute to overall bouquet and to fruit, floral, and herbal sensory descriptors. In grape, monoterpenes are synthesized via the methylerythritol phosphate (MEP) pathway, with 1-deoxy-D-xylulose-5-phosphate synthase (DXS) catalyzing the first committed step (Dunlevy *et al*., 2009). Several terpenoid-associated QTLs have been identified in *V. v. vinifera*, including a locus on chromosome 5 containing *VviDXS* associated with increased monoterpenoid accumulation in Muscat cultivars and aromatic mutants of otherwise neutral backgrounds (Duchêne *et al*., 2009a,b). Structural diversity among monoterpenes is generated by terpene synthases (TPSs), which form a large, often tandemly duplicated gene family in grapevine and show substantial cultivar-specific variation (Martin *et al*., 2010; Drew *et al*., 2016; Smit *et al*., 2021; Bosman *et al*., 2023). However, it remains unclear whether variation in monoterpenoid accumulation in moderately aromatic cultivars is primarily driven by structural variation at terpene biosynthetic loci, differences in gene dosage, or regulatory variation affecting gene expression.

Previous studies of monoterpenoid production in *V. v. vinifera* have focused largely on highly aromatic cultivars such as Muscat blanc and its derivatives, whereas the genetic basis of moderate monoterpenoid accumulation in cultivars such as Riesling and Viognier remains poorly characterized (Lin *et al*., 2019). Most analyses of grape volatile composition have relied on interspecific hybrids or table grape crosses, often with limited genetic resolution and without genome references, constraining the identification of causal genes and regulatory mechanisms. In contrast, relatively few studies have examined aroma variation in intraspecific wine grape populations. This gap is notable given the genetic divergence between wine and table grapes, the greater contribution of *V. v. sylvestris* ancestry in wine grapes, and the structural complexity of grapevine genomes (Aradhya *et al*., 2003; Arroyo-García *et al*., 2006; Zhou *et al*., 2017, 2019). Together, these limitations have hindered a mechanistic understanding of the genetic architecture of aroma traits in wine grapes.

Riesling and Cabernet Sauvignon represent contrasting aroma profiles within *V. v. vinifera*. Riesling is defined by moderate monoterpenoid levels and a strong contribution of norisoprenoids such as 1,1,6-trimethyl-1,2-dihydronaphthalene (TDN) to its varietal profile (Strauss *et al*., 1987; Marais and Rapp, 1991; Kalua and Boss, 2010), whereas Cabernet Sauvignon accumulates low levels of monoterpenes and is characterized by methoxypyrazines (Kalua and Boss, 2010; Dunlevy *et al*., 2013). Both cultivars are widely planted, genetically representative of Western European germplasm, and extensively used in breeding, making them well-suited for dissecting the genetic basis of aroma variation (Alston and Sambucci, 2019).

To dissect the genetic basis of aroma variation in wine grape, we present a chromosome-scale diploid genome assembly of Riesling and use it to anchor genotyping-by-sequencing and high-density linkage mapping in an F1 Riesling × Cabernet Sauvignon population. Coupling this framework with semi-targeted metabolomics of free and glycosidically bound volatiles, we map QTLs associated with a broad spectrum of berry aroma compounds. Integration with haplotype-resolved expression data identified two major loci controlling (3*S*)-linalool accumulation, with haplotype-specific expression of *VviTPS54* emerging as a key determinant of monoterpenoid variation between the parental cultivars and their progeny.

## Materials and Methods

### RxCS Mapping Population

The population used in this study consisted of 138 full-sibling progeny resulting from a cross between *V. v. vinifera* cv. Riesling clone FPS 24/GM 110 and *V. v. vinifera* cv. Cabernet Sauvignon clone FPS 08 planted at the Viticulture and Enology Department Experimental Station (Oakville, Napa, CA) in 2018. This planting was previously used for identifying QTLs related to microbiome recruitment, cluster architecture, and yield components (Flörl *et al*., 2025; Sharma *et al*., 2025). The population was planted as a randomized complete block vineyard consisting of three biological replicates of each genotype in three adjacent blocks for a total of 9 vines per genotype, including both parent varieties. The experimental vines were completely surrounded by a buffer of Cabernet Sauvignon vines to minimize edge effects. Rows were oriented northwest-southeast with 2.4 meters between rows and 1.8 meters between vines. Training consisted of bilateral cordons in a modified vertical shoot positioning system pruned to two-bud spurs. Water was supplied through drip irrigation. Pesticide applications and irrigation were scheduled at the discretion of the vineyard manager.

### Genome Assembly and Annotation

The genome assembly of Riesling was scaffolded at chromosome level using contigs generated previously (Zou *et al*., 2021). Using a high-density consensus map and HaploSync (Zou *et al*., 2020; Minio *et al*., 2022), the contigs were phased and anchored into 19 sets of diploid chromosomes after 7 runs of HaploSplit and 4 runs of HaploFill. Gene models were functionally annotated concatenating the results from blastp v.2.7.1+ (Camacho *et al*., 2009) and interproscan v.5.27-66.0 (Jones *et al*., 2014b) using blast2go (Gotz *et al*., 2008) as described previously (Vondras *et al*., 2019). The parentally phased Cabernet Sauvignon genome assembly and annotation were obtained from (Cochetel *et al*., 2025).

### Genotyping, Map Construction, and QTL Analysis

A consensus linkage map was generated from the Riesling × Cabernet Sauvignon (RxCS) genotyping-by-sequencing (GBS) data previously obtained by Flörl et al. (2025). GBS data were aligned to the Riesling FPS24 Hap1 genome sequence using BWA-MEM (ver. 0.7.17-r1188) with default parameters (Li and Durbin, 2009). SNPs were filtered with Tassel 5 using the following criteria: a coverage depth of >6, minor allele frequency of >0.1, and missing data <10% (Bradbury *et al*., 2007). SNPs within 64 base pairs were merged. Map construction followed the method published in Flörl et al. (2025) and Lopez-Moreno *et al*. (2023) using the Kosambi mapping function in the ASMap package (Taylor and Butler, 2017). Collinearity between the genetic map and physical map was assessed by calculating the Spearman correlation coefficient. Mapchart v.2.2 was used to draw linkage groups and QTL intervals (Voorrips, 2002). QTL analysis was performed using MapQTL v.6 (Van Ooijen, n.d.) on genotype means. The maximum likelihood mixture model was used to locate QTLs and permutation tests of 1000 permutations were used to identify genome-wide significance thresholds at 0.05 significance level. The physical boundaries of significant QTLs were located on the haplotype 1 of the Riesling genome using the coordinates of two flanking markers roughly 1-2 LOD from the peak (Lander and Botstein, 1989).

### Sample Collection and Processing

Berries from the RxCS population were collected for volatile and glycoside analysis from 128 genotypes in 2020 and from a subset of 8 genotypes in 2021. Berries were sampled from the two parent cultivars in both years. For the 2020 data set, berries were sampled once or twice a week starting 11 weeks post-anthesis to track sugar accumulation. Collection for chemical analysis occurred when the average Brix reached 23°Brix or when sugar levels plateaued for three consecutive samplings. Approximately 150g of berries were collected by snipping off individual berries from multiple clusters. Each block replicate consisted of pooled berries from the three replicate vines of each genotype. Berries were frozen in liquid nitrogen and stored at -80° until further processing. The 2021 samples were collected in the same manner at two time points: 14 weeks post anthesis (Preharvest) and at full ripeness as determined with the same methodology as in 2020 (Harvest). Homogenized frozen tissue for HS-SPME-GC-MS and UHPLC-QTOF/MS analysis was prepared according to the protocol described by (Hjelmeland *et al*., 2015). Approximately 60g of whole frozen berries with pedicels removed were weighed and cooled in liquid nitrogen to facilitate grinding. An A11 Basic Grinder (IKA, Staufen, Germany) was used to homogenize the berries to a fine powder. Subsequently, the powder was transferred into 50mL Falcon tubes and refrozen in liquid nitrogen. Homogenized samples were then stored at -80°C until extraction. Samples were collected for RNA extraction in 2021 from the eight progeny genotypes and both parent cultivars concurrently with collection for metabolite analysis. Three replicates of each genotype were collected, one per block. Each sample consisted of a pool of 30 berries collected from three vines. Berries were kept cool during transport. Pedicels and seeds were removed and samples frozen in liquid nitrogen before storage at -80°C until extraction.

### Headspace solid-phase microextraction coupled with gas chromatography-mass spectrometry (HS-SPME-GC-MS) analysis

Volatile compounds were analyzed semi-quantitatively using the HS-SPME-GC-MS protocol used by (Hendrickson *et al*., 2016). Analysis of all samples used an Agilent 6890N gas chromatograph coupled to a 5975 mass selective detector (Agilent, Santa Clara, CA, USA) with an MPS2 autosampler (Gerstel, Mülheim an der Ruhr, Germany). Berry samples were analyzed in duplicate for a total of 6 samples per genotype (2 technical replicates of the 3 block replicates). For each sample, 20 (± 0.20) g of frozen homogenized powder were measured into 50mL polypropylene centrifuge tubes with 5 mL of 1.0 M sodium citrate adjusted with HCl to pH 6. The mixture was homogenized and allowed to thaw completely before centrifuging for 15 min at 4°C and 5,000 rpm. Two 10mL aliquots of supernatant were pipetted into 20 mL amber headspace vials with 5 mg of NaCl and spiked with 50μl each of 10 mg/L 2-octanol and 2-undecanone dissolved in 100% ethanol as internal standards. All samples were immediately capped with 20 mm crimp caps with PTFE/silicone septa (Supelco, Bellefonte, PA, USA) and analyzed within 24 hours of extraction. Sample order was randomized for each run.

The MPS2 sampling sequence was programmed so that berry samples were equilibrated at 30°C for 5 min with agitation in the MPS2 incubator prior to extraction using a 23-gauge syringe, 1 cm polydimethylsiloxane (PDMS) SPME fiber for 45 minutes at 30°C and 250 rpm. The fiber was thermally desorbed in the GC inlet at 260°C operating in splitless mode. Chromatographic separation utilized a DB-Wax ETR capillary column (30 m, 0.25 mm i.d., 0.25 μm film thickness) with helium as the carrier gas at a constant flow of 0.90 mL.min⁻¹. The oven temperature program began at 40°C for 5 min, ramped at 3°C.min⁻¹ to 180°C, then 30°C.min⁻¹ to 260°C with a final hold of 10 min. The MS transfer line, ion source, and quadrupole were maintained at 260°C, 230°C, and 150°C, respectively. Mass spectra were acquired in scan mode over an *m/z* range of 30–300 with a 2 min solvent delay. Compounds were identified using NIST mass spectrum matches and Kovats retention indices (RI). Analysis of GCMS data was performed on Agilent MassHunter Qualitative and Quantitative packages (versions 10.0). Relative abundances were calculated by normalizing compound peak areas to the peak area of the 2-octanol internal standard.

### Solid-Phase-Extraction (SPE) of Monoterpene Glycosides

Samples for glycoside analysis were extracted according to the solid-phase extraction (SPE) method used by Hjelmeland et al. (Hjelmeland *et al*., 2015). Five g of previously prepared homogenized berry powder were added to 5 mL of 50 mM sodium citrate (pH 5) and vortexed. After thawing, samples were centrifuged at 4°C and 4100 x *g* for 15 minutes. 5 mL of supernatant were transferred to clean Falcon tubes with 50 μl of 10,000 mg/L decyl-β-D-glucopyranoside internal standard. Solid phase extractions were performed in 96-well C18 phase Agilent plates with 100 mg bed mass per plate using a 96-well acrylic vacuum manifold. The plates were conditioned with methanol and 18 MΩ water, and the samples were washed and eluted as outlined with 1 mL of methanol. Samples were then stored at -80°C until analysis. Analysis of glycoside samples occurred no more than two weeks after extraction.

### UHPLC-QTOF/MS Analysis

UHPLC-QTOF/MS analysis was conducted on an Agilent 1290 UHPLC coupled to an Agilent 6545 QTOF mass spectrometer with an Agilent Dual ESI Jet Stream source in negative ion mode. Separation was achieved using an Agilent Poroshell 120 phenyl-hexyl column (2.1x 150 mm, 2.7 μm) with an Agilent Poroshell 120 phenyl-hexyl (2.1 x 5 mm, 2.7 μm) guard column at 40°C. Mobile phases consisted of water with 0.1% acetic acid (A) and acetonitrile with 0.1% acetic acid (B) delivered at 0.42 mL.min⁻¹. The gradient program increased from 5% B to 20% over 5 minutes, held at 20% B for 13 minutes, then ramped up to 90% B over 4 minutes before returning to initial conditions for equilibration of the column. Samples were injected at a volume of 9 µL. Mass spectra were acquired in negative ion mode over an *m/z* range of 100–1000 using the Agilent “All ions” workflow and collision energies of 15 and 30 V. Continuous reference mass correction using proton abstracted purine (*m/z* = 119.0362) and hexakis acetate adduct (*m/z* = 980.016375) was applied to maintain mass accuracy. Target compounds were extracted using a ± 20 ppm mass window and retention time tolerance of ± 0.5 min. Monitoring of instrument performance was done regularly using a control sample (E&J Gallo Brandy (Modesto, CA, USA)). Glycoside data were normalized to decyl-β-D-glucopyranoside equivalents prior to analysis and were therefore semiquantitative.

### RNA extraction, sequencing, and analysis

Samples for RNA extraction were processed on the collection day as described in Blanco-Ulate et al. (Blanco-Ulate *et al*., 2013, 2017). Briefly, deseeded berries were frozen using liquid nitrogen and ground with a mechanical mill. Two g of tissue were used for RNA extraction. Total RNA extraction was performed on 60 samples (10 genotypes at two timepoints and three biological replicates). Extraction and library preparation, were performed as in (Vondras *et al*., 2021). RNA-seq libraries were sequenced in 75 single-end mode, using an Illumina NextSeq sequencer (DNA Technology Core Facility, University of California, Davis).

### Transcript abundance quantification

The pipeline nf-core/rnaseq v.3.13.2 was used to perform gene expression analysis using default parameters (Ewels *et al*., 2020; Harshil Patel *et al*., 2026). Briefly, raw RNA-Seq reads were quality-filtered using TrimGalore (https://github.com/FelixKrueger/TrimGalore). Filtered reads were then pseudo-aligned using Salmon (Patro *et al*., 2017). The output salmon.merged.gene_tpm.tsv was used for the subsequent analyses. To aid in the identification of genes related to aroma compound metabolism, differential expression analysis was carried out on samples from the parent cultivars at both sampling timepoints using the R package DESeq2 v.1.42.1 (Love *et al*., 2014). Protein sequence motifs and domains were identified using TargetP 2.0 (https://services.healthtech.dtu.dk/services/TargetP-2.0/) and InterPro 108.0 (https://www.ebi.ac.uk/interpro/) (Jones *et al*., 2014a; Almagro Armenteros *et al*., 2019; Blum *et al*., 2025).

### Statistical Analysis

Multivariate analysis of variance (MANOVA) and univariate analysis of variance (ANOVA) were used to test for statistical significance for all HS-SPME-GC-MS and UHPLC-QTOF/MS data with a significance threshold of 0.05. Correlations between metabolite and RNA-Seq data were assessed through Spearman’s rank-order correlation using *cor.test* in R Statistical Software (v4.1.2, R Core Team 2021). Significance of correlation was determined by Benjamini-Hochberg adjusted *P*-value calculated using the *corTest* function from the R package psych v 2.5.6 (Revelle, 2026). Multivariate ordination analyses were carried out using the FactoMineR v. 2.13 and factoextra v. 1.0.7 packages (Lê *et al*., 2008; Kassambara and Mundt, 2016). Partial least squares discriminant analysis (PLS-DA) of parental metabolites was conducted on Metaboanalyst 6.0 (https://www.metaboanalyst.ca/home.xhtml) (Pang *et al*., 2024).

## Results

### Segregation of volatile compounds within the Riesling × Cabernet Sauvignon population

Ripe berries from Riesling, Cabernet Sauvignon, and their F1 progeny were analyzed by HS-SPME-GC-MS to characterize volatile composition. Two genotypes did not produce fruit, and an additional 10 genotypes yielded insufficient material for analysis. In total, 186 volatile compounds were detected, of which 110 differed significantly among genotypes. Aldehydes and alcohols represented the largest fraction of detected compounds (**Supplementary Tables S1 and S2**; **Supplementary Fig. 1**). Additional classes included acids, alkanes, aromatic compounds, esters, furans, ketones, mono- and sesquiterpenes, norisoprenoids, and a small group of miscellaneous compounds. Overall, alcohols and aldehydes dominated the volatile profile.

Riesling and Cabernet Sauvignon showed clear differences in volatile composition. Partial least squares discriminant analysis (PLS-DA) identified major discriminating compounds, including C6 and C7 alcohols and related metabolites, mono- and sesquiterpenes, one norisoprenoid, two acetate esters, and two phenylpropanoid-derived compounds (**Fig. 1A-B**). These largely overlap with the compounds previously reported to distinguish the two cultivars (Kalua and Boss, 2010). Although (3*S*)-linalool was not detected in that study, Riesling was distinguished by higher monoterpene content, a pattern confirmed here. In the present dataset, total monoterpenes were substantially higher in Riesling (**Fig. 1C**), and composition also differed between cultivars, with (3*S*)-linalool predominating in Riesling and geraniol in Cabernet Sauvignon (**Supplementary Table S1**).

**Figure 1.**
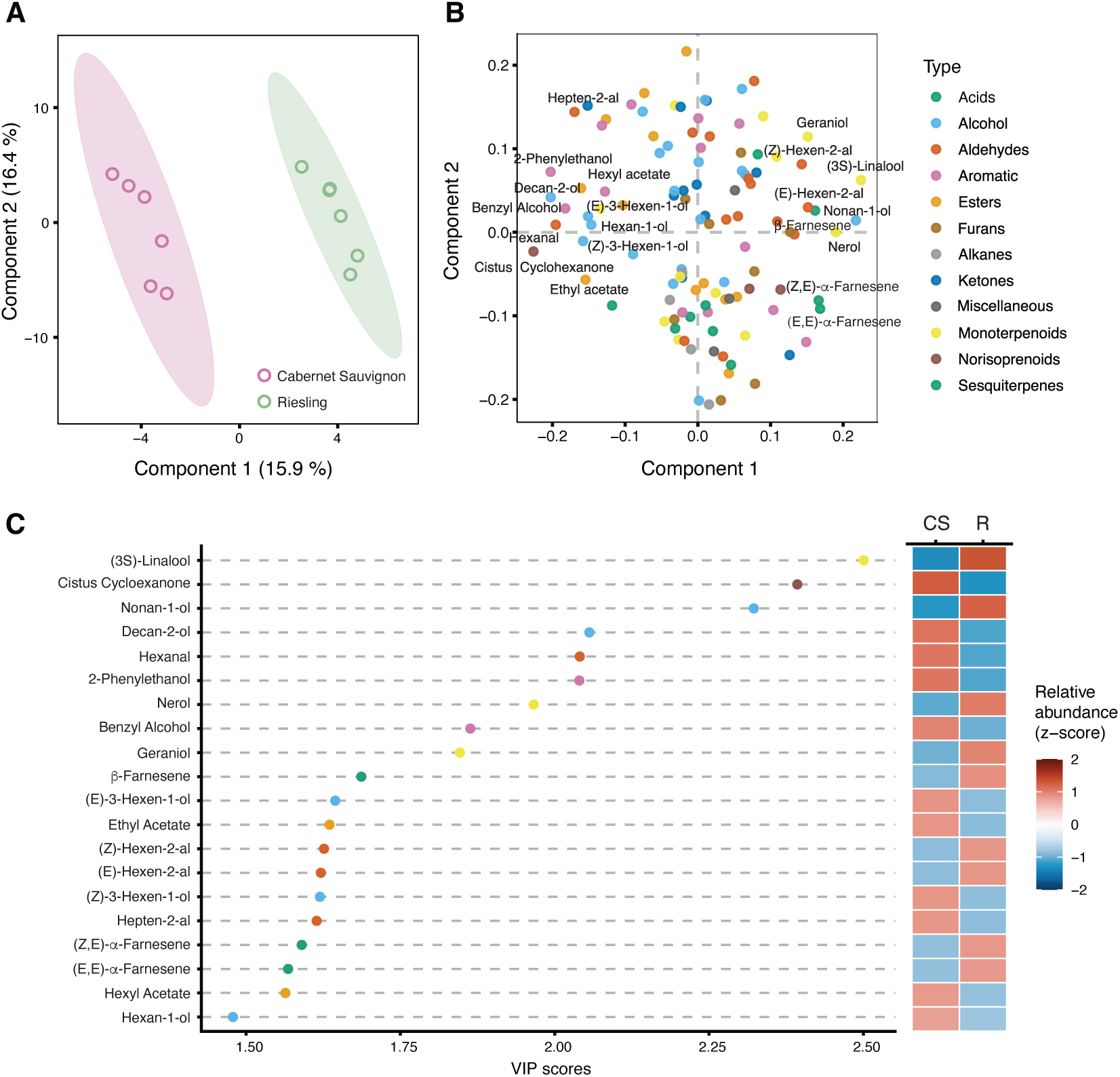
Partial least squares discriminant analysis (PLS-DA) of volatile compounds in Riesling and Cabernet Sauvignon berries. (A) PLS-DA score plot showing separation between cultivars. Points represent individual samples; ellipses indicate 95% confidence intervals for each group. (B) Loading plot showing the contribution of individual volatile compounds to sample separation. Points are colored by compound class. (C) Variable importance in projection (VIP) scores for the top 20 compounds contributing to cultivar discrimination. Points are colored by compound class. The accompanying heatmap shows scaled relative abundance (z-score) of each compound in Cabernet Sauvignon (CS) and Riesling (R).

Hierarchical clustering by Euclidean distance and complete linkage of metabolite relative abundance across genotypes yielded nine clusters (**Supplementary Table S1**). Clusters broadly reflected biosynthetic relationships, with lipoxygenase-derived compounds grouping together (clusters A, B, and E) and terpenoids forming distinct clusters, including a monoterpene-rich cluster (cluster I) containing (3*S*)-linalool, geraniol, and nerol. Sesquiterpenes and norisoprenoids co-clustered (clusters D and F), whereas remaining clusters were more heterogeneous but still showed partial organization by chemical structure. Overall, the population exhibited broad variation in metabolite abundance, with many genotypes showing strongly positive or negative Z-scores across clusters. These patterns indicate segregation of volatile traits within the population and suggest that subsets of metabolites are subject to shared genetic control.

The abundance of representative compounds from each cluster across parents and progeny is shown in **Fig. 2**. Several compounds, including linalool, exhibited transgressive segregation within the mapping population. Notably, (3*S*)-linalool (cluster I) showed clear differentiation between parental genotypes and extensive segregation in the progeny, with some individuals exceeding Riesling levels. Broad-sense heritability (H²) was estimated for each significant compound (**Supplementary Table S2**), ranging from 0.15 for β-myrcene to 0.96 for (*Z*)-theaspirane. Monoterpenes showed consistently high heritability, with H² values of 0.90 for (3*S*)-linalool and geraniol, and 0.73 for nerol, indicating strong genetic control of monoterpenoid biosynthesis.

**Figure 2.**
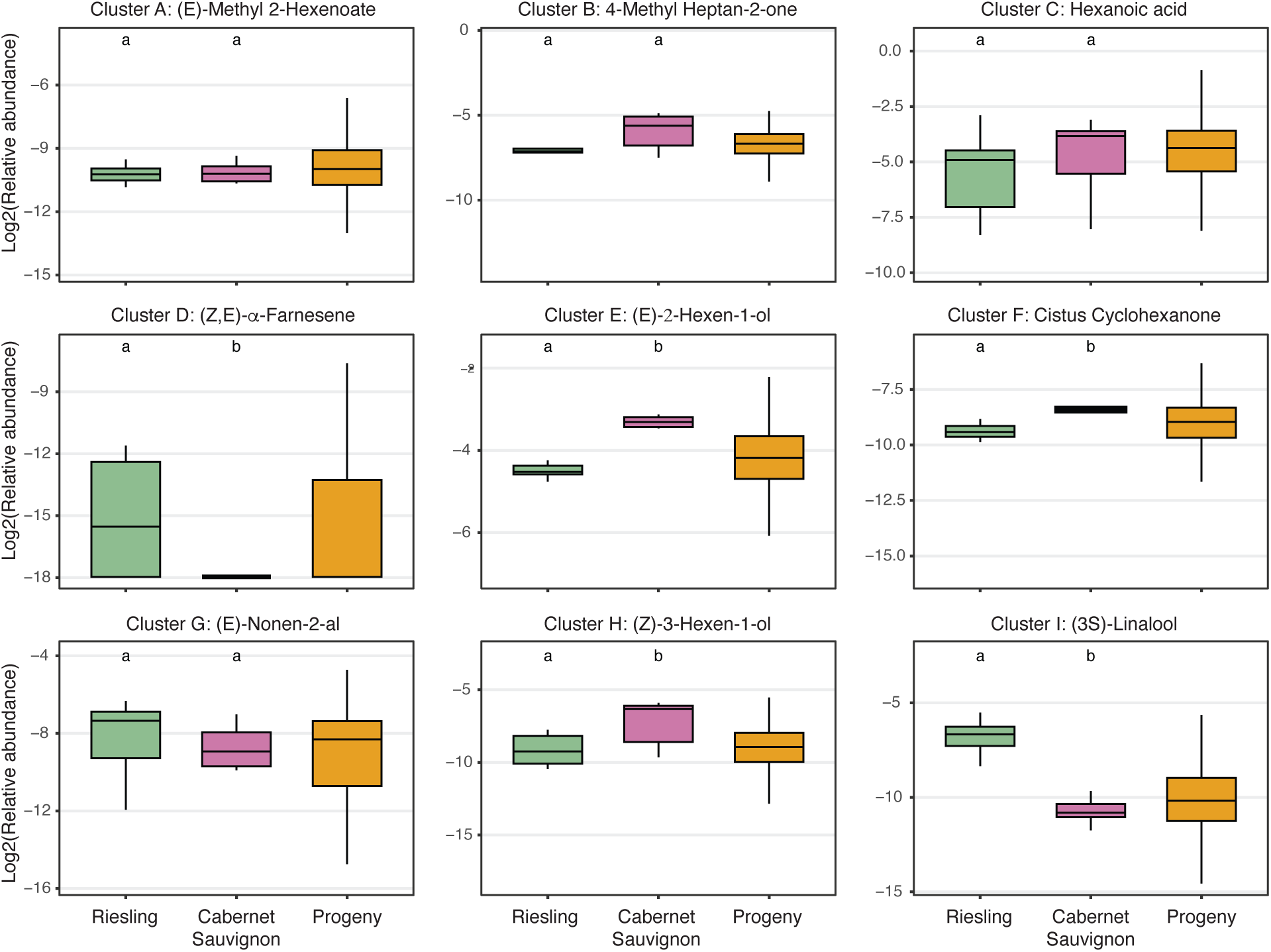
Distribution of representative volatile compounds from hierarchical clusters in Riesling, Cabernet Sauvignon, and their F1 progeny. Boxplots show log_2_-transformed relative abundance of representative compounds from clusters A-I identified by hierarchical clustering based on compound abundance across genotypes. Colors indicate genotype groups (Riesling, Cabernet Sauvignon, and progeny). Titles denote compound identity and cluster assignment; compound classes are indicated by panel titles. Letters above boxplots represent Tukey’s HSD groupings (*P* < 0.05).

### Segregation of monoterpene glycosides within the RxCS population

Because most monoterpenes in grape occur as nonvolatile glycosides (Hjelmeland and Ebeler, 2015), we quantified the relative abundance of 14 monoterpenoid glycosides in the RxCS population (**Table 1**). Monoterpene glycosides were classified as pentose-hexose conjugates (PH; PentHex-MT-ol) and deoxyhexose-containing conjugates (DH; DeoxyHex-MT-ol).

**Table 1.** Relative abundance of monoterpene glycosides. Mean relative abundances in decyl-β-D-glucopyranoside equivalents are shown for 12 pentose-hexose monoterpenol glycosides (PH) and two deoxyhexose monoterpenol glycosides (DH). RT, retention time (min); R and CS, mean relative abundance in Riesling and Cabernet Sauvignon parents, respectively; Progeny, mean relative abundance across the RxCS population; Min and Max, minimum and maximum relative abundance observed among the progeny.

| Monoterpenol Glycoside | RT | R Mean | CS Mean | Progeny Mean | Min | Max |
| --- | --- | --- | --- | --- | --- | --- |
| PentHex-MT-ol 1 (PH1) | 8.97 | 2.02 | 0.186 | 0.925 | 0.0133 | 7.868 |
| PentHex-MT-ol 2 (PH2) | 9.71 | 0.84 | 0.136 | 0.853 | 0.0059 | 8.936 |
| PentHex-MT-ol 3 (PH3) | 10.38 | 0.16 | 0.020 | 0.077 | 0.0010 | 0.584 |
| PentHex-MT-ol 4 (PH4) | 10.48 | 0.32 | 0.020 | 0.086 | 0.0016 | 1.471 |
| PentHex-MT-ol 5 (PH5) | 10.90 | 4.14 | 0.001 | 0.685 | 0.0010 | 13.530 |
| PentHex-MT-ol 6 (PH6) | 11.59 | 0.43 | 0.002 | 0.060 | 0.0003 | 1.050 |
| PentHex-MT-ol 7 (PH7) | 11.90 | 0.82 | 0.000 | 0.087 | 0.0002 | 2.504 |
| PentHex-MT-ol 8 (PH8) | 12.18 | 1.14 | 0.066 | 0.277 | 0.0628 | 1.594 |
| DeoxyHex-MT-ol 1 (DH1) | 12.23 | 2.24 | 0.000 | 0.296 | 0.0001 | 7.157 |
| PentHex-MT-ol 9 (PH9) | 12.72 | 0.68 | 0.003 | 0.069 | 0.0034 | 1.312 |
| PentHex-MT-ol 10 (PH10) | 13.39 | 2.60 | 0.275 | 1.075 | 0.1443 | 5.860 |
| PentHex-MT-ol 11 (PH11) | 14.06 | 0.24 | 0.035 | 0.072 | 0.0021 | 0.297 |
| PentHex-MT-ol 12 (PH12) | 14.36 | 0.31 | 0.030 | 0.105 | 0.0007 | 0.739 |
| DeoxyHex-MT-ol 2 (DH2) | 14.46 | 0.15 | 0.002 | 0.031 | 0.0004 | 0.307 |

Principal component analysis (PCA) revealed substantial variation in glycoside accumulation, with the first two components explaining 66.3% of total variance (**Fig. 3**). Riesling and Cabernet Sauvignon were clearly separated along PC1, while progeny were distributed between parental extremes. PC1 primarily reflected overall monoterpene glycoside abundance, driven largely by pentose-hexose conjugates PH5-PH9. In contrast, separation along PC2 was associated with glycoside composition, with PH1-PH4 showing strong positive correlations and contributing most to this axis. The correlations between monoterpene glycosides and free monoterpenoids are shown in **Fig. 4**. Most glycosides were significantly correlated with at least one aglycone (adjusted *P* < 0.05), although correlations were generally modest (r < 0.7). Geraniol showed weak associations, limited to PH10 and PH12. In contrast, (3*S*)-linalool exhibited strong correlations (r > 0.8) with PH5, PH7, and DH1, indicating tight coupling between glycoside pools and free linalool levels. Notably, PH1 and PH2 showed little to no correlation with any detected monoterpenoids despite their relatively high abundance in some genotypes.

**Figure 3.**
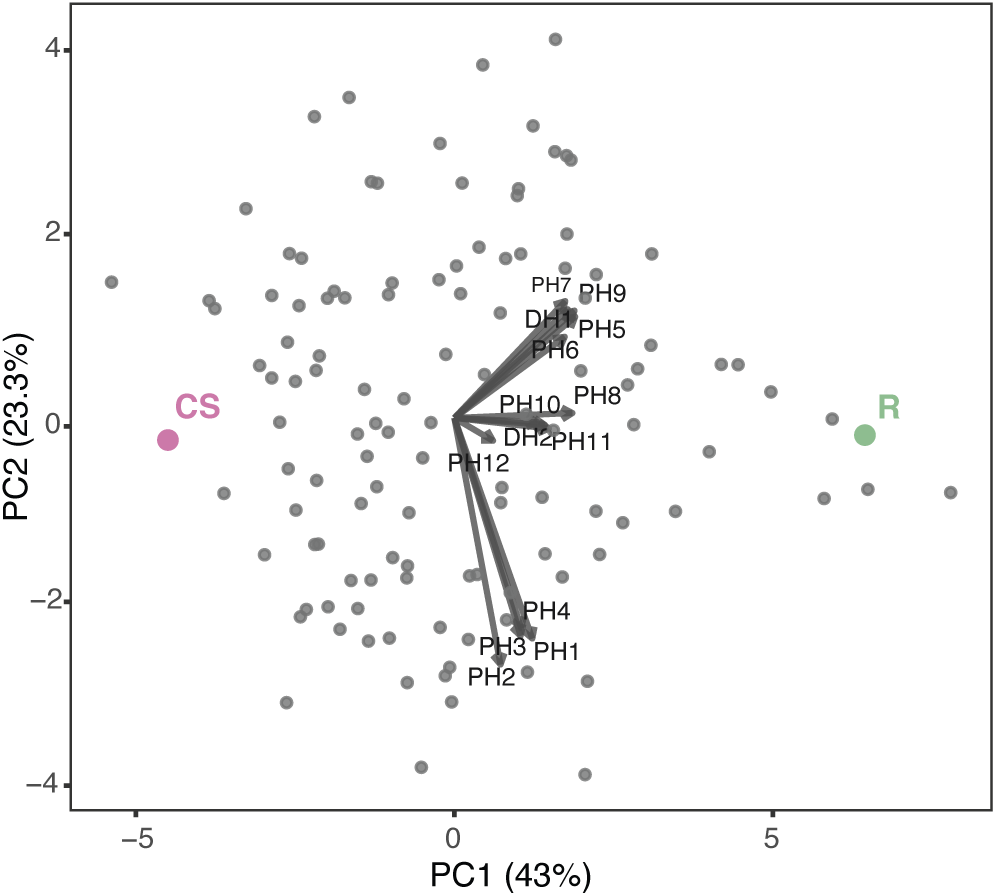
Principal component analysis of monoterpene glycosides in the RxCS population. Genotype means of individual progeny are shown as gray points, with parental genotypes highlighted (Riesling, R; Cabernet Sauvignon, CS). The first two principal components explain 66.3% of total variance (PC1: 43.0%; PC2: 23.3%). Arrows indicate loadings of individual glycosides (PH and DH), with direction and length reflecting their contribution to each principal component. PC1 primarily captures variation in total glycoside abundance, whereas PC2 reflects differences in glycoside composition.

**Figure 4.**
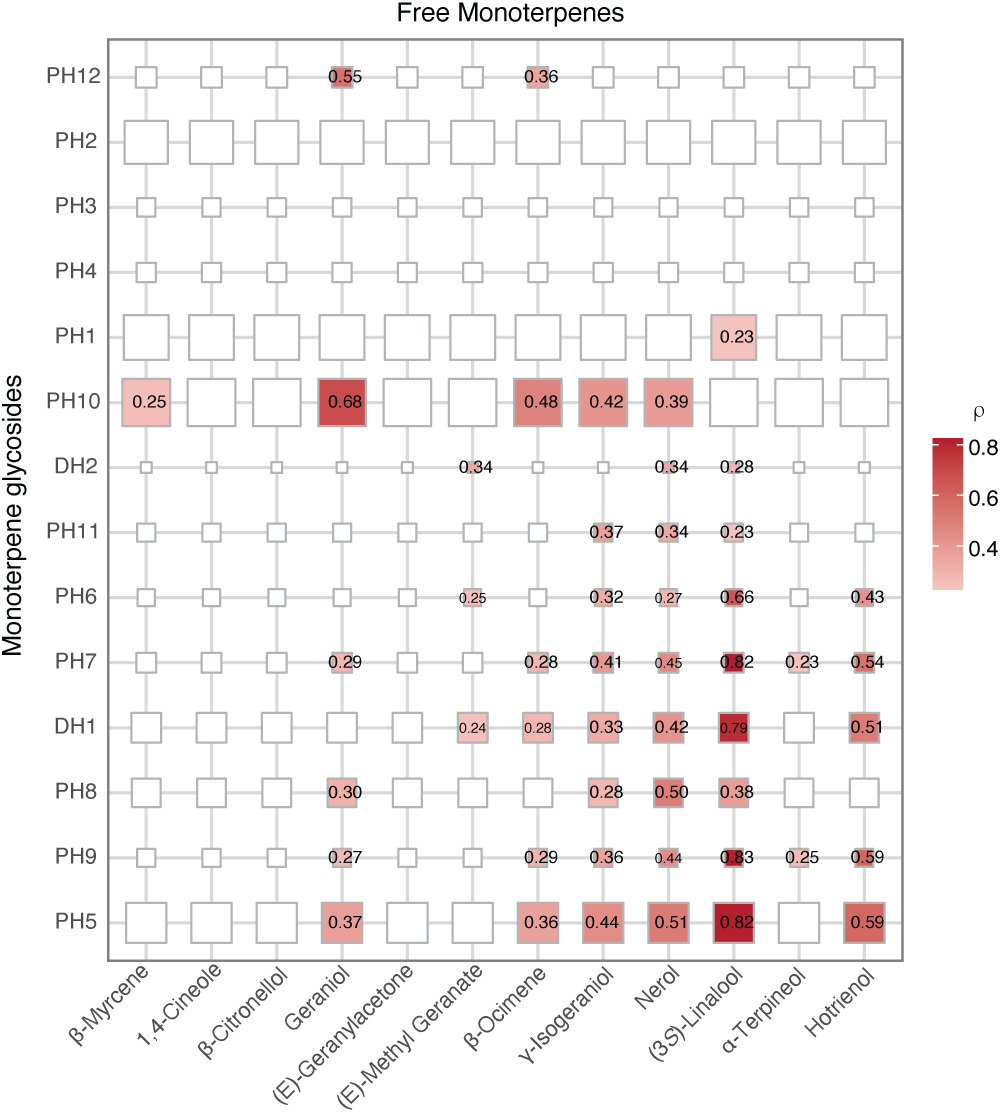
Correlation between monoterpene glycosides and volatile monoterpenes in the RxCS population. Heatmap showing pairwise Spearman’s rank correlation coefficients (ρ) between monoterpene glycosides (rows) and free monoterpenes (columns). Color intensity reflects the strength of positive correlations, and values are shown within each cell. Only significant correlations (adjusted *P* < 0.05) are displayed. Rows are ordered based on PCA structure (PC1 and PC2) shown in Fig. 3. Tile size is proportional to the mean relative abundance of each glycoside in the population.

### Assembly and annotation of the Riesling genome and linkage map construction

The genome of *Vitis vinifera* cv. Riesling clone FPS24 was assembled to chromosome scale using HaploSync (Minio *et al*., 2022) in combination with a high-density consensus map (Zou *et al*., 2020). The two phased haplotypes (Haplotype 1, RHap1; Haplotype 2, RHap2) were comparable in size (**Table 2**), each consisting of ∼48.4% repetitive sequence and over 31,000 annotated protein-coding gene loci. Unplaced sequences were more repeat-rich (57.4%) and accounted for fewer than 10% of the annotated genes.

**Table 2.** Summary statistics of the diploid genome assembly of Riesling.

|  | Haplotype 1<br>(RHap1) | Haplotype 2<br>(RHap2) | Unplaced |
| --- | --- | --- | --- |
| Sequence length (Mb) | 489.62 | 494.15 | 108.17 |
| Number of sequences | 19 | 19 | 1,842 |
| N50 (bp) | 27,222,099 | 27,741,044 | 66,238 |
| Repeats (%) | 48.48 | 48.45 | 57.37 |
| Genes | 31,292 | 31,521 | 6,049 |

To enable QTL mapping of aroma traits in the segregating population, a genetic linkage map was constructed using GBS data. A total of 134,411 100-bp reads were obtained, with a mean read depth of 15.1x (median 2.9x; SD 90.2). After filtering for read depth, missing data, and minor allele frequency, 7,680 high-quality SNPs were retained for map construction. A consensus parental map was generated for QTL analysis of aroma compounds (**Fig. 5**). The final linkage map comprised 5,186 markers across 19 linkage groups, spanning 1,346.36 cM with an average inter-marker distance of 0.26 cM. The largest gap between markers was 13.56 cM (**Supplementary Table S3**). The genetic map showed high collinearity with the physical map (**Supplementary Fig. S2**), with an average Spearman correlation coefficient of 0.98.

**Figure 5.**
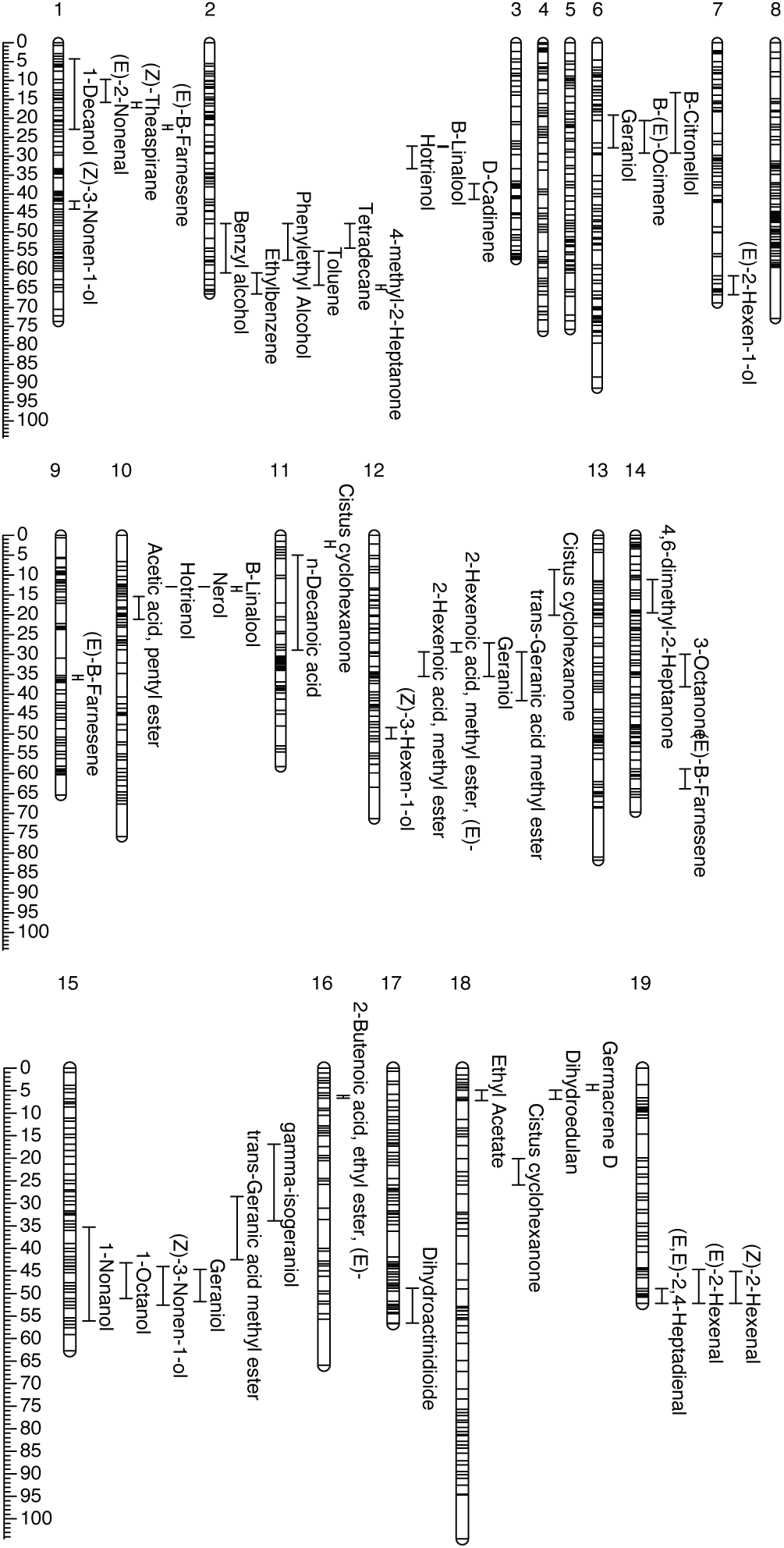
Linkage map and QTLs associated with volatile aroma compounds. Genetic linkage map of the Riesling × Cabernet Sauvignon F1 population showing 19 linkage groups corresponding to the 19 grapevine chromosomes. Marker density is represented along each linkage group, with positions in centimorgans (cM). Horizontal bars indicate the positions of QTLs associated with individual volatile compounds, with labels denoting compound identity. QTL intervals correspond to confidence intervals defined from the mapping analysis. The scale bar (left) indicates genetic distance (cM).

### Detection of QTLs associated with volatile and bound aroma compounds

QTL analysis identified genomic regions associated with variation in aroma compound abundance (**Fig. 5**). A total of 40 volatile compounds were associated with significant QTLs, with individual loci explaining 16.4% (2-phenylethanol) to 65.5% (δ-cadinene) of phenotypic variance, both on chromosome (Chr) 2. Most compounds (33) were associated with a single QTL, while seven showed multiple QTLs, for a total of 49 loci distributed across the genome. No significant QTLs were detected on Chr 3, 4, 5, 8, or 13, whereas Chr 2 harbored the highest number (nine loci). Monoterpenes accounted for the largest number of QTLs (13 loci). No significant QTLs were detected for furans or miscellaneous compounds. Compounds with significant QTLs are listed in **Table 3**, together with their physical positions in the RHap1 assembly. QTLs frequently colocalized for compounds sharing structural or biosynthetic relationships. For example, the aliphatic alcohols octan-1-ol, nonan-1-ol, and (*Z*)-3-nonen-1-ol mapped to the same region on Chr 15, while phenylalanine-derived volatiles (benzyl alcohol and 2-phenylethanol) colocalized on Chr 2.

**Table 3.** Significant quantitative trait loci (QTLs) associated with volatile compounds and monoterpene glycosides in the RxCS population. QTL intervals were defined by the positions of flanking markers corresponding to a 0.5-LOD drop on either side of the peak. Only QTLs exceeding the 95th percentile of the genome-wide LOD threshold are shown.

| Class | QTL Number | Compound | Chr | Start (bp) | End (bp) | Start (cM) | End (cM) | Peak LOD | Peak Marker (RHap1) | Percent Variance | Genes in QTL |
| --- | --- | --- | --- | --- | --- | --- | --- | --- | --- | --- | --- |
| Acid | 1 | n-Decanoic acid | 11 | 3,048,855 | 5,902,730 | 5.04 | 28.91 | 4.99 | CHR11_3519187 | 17 | 290 |
| Alcohol | 2 | (*E*)-2-hexen-1-ol | 1 | 694,440 | 5,665,110 | 4.29 | 22.89 | 5.63 | CHR01_2204154 | 19 | 496 |
|  | 3 | (*Z*)-3-hexen-1-ol | 15 | 13,488,830 | 17,681,341 | 35.31 | 56.13 | 8.15 | CHR15_16690679 | 26.3 | 404 |
|  | 4 | Nonan-1-ol | 15 | 15,004,444 | 16,946,524 | 43.22 | 51.13 | 9.7 | CHR15_15875293 | 30.5 | 201 |
|  | 5 | Octan-1-ol | 7 | 26,315,802 | 27,899,218 | 61.56 | 66.56 | 5.56 | CHR07_29254069 | 18.8 | 174 |
|  | 6 | (*Z*)-3-nonen-1-ol | 12 | 19,238,044 | 21,186,879 | 48.35 | 51.2 | 8.35 | CHR12_20514952 | 26.9 | 133 |
|  | 7 | (*Z*)-3-nonen-1-ol | 1 | 14,963,421 | 18,073,634 | 41.85 | 43.99 | 5.89 | CHR01_14905497 | 19.8 | 92 |
|  | 8 | Decan-1-ol | 15 | 15,570,482 | 17,009,996 | 43.95 | 52.55 | 7.4 | CHR15_16157018 | 24.2 | 152 |
| Aldehyde | 9 | (*E,E*)-2,4-heptadienal | 19 | 19,881,449 | 21,925,778 | 48.87 | 52.2 | 7.93 | CHR19_20030878 | 25.7 | 134 |
|  | 10 | (*E*)-hexen-2-al | 19 | 15,804,112 | 21,733,333 | 44.69 | 52.2 | 6.88 | CHR19_20748095 | 22.7 | 286 |
|  | 11 | (*Z*)-hexen-2-al | 19 | 17,399,164 | 21,925,778 | 45.05 | 52.2 | 5.14 | CHR19_20030878 | 17.5 | 242 |
|  | 12 | (*E*)-nonen-2-al | 1 | 2,676,209 | 3,653,671 | 9.65 | 15.75 | 8.68 | CHR01_2720122 | 27.8 | 100 |
| Aromatic | 13 | Ethylbenzene | 2 | 11,073,778 | 19,214,008 | 47.83 | 60.84 | 5.53 | CHR02_15537002 | 18.7 | 284 |
|  | 14 | Toluene | 2 | 19,214,008 | 21,275,965 | 60.84 | 66.36 | 5.8 | CHR02_20358829 | 19.5 | 110 |
|  | 15 | Benzyl alcohol | 2 | 11,566,036 | 19,138,182 | 47.83 | 57.52 | 4.8 | CHR02_15851855 | 16.4 | 265 |
|  | 16 | 2-Phenylethanol | 2 | 14,128,650 | 21,083,470 | 55.1 | 64.07 | 6.52 | CHR02_20358829 | 21.7 | 287 |
| Ester | 17 | Methyl hex-2-enoate | 16 | 4,593,278 | 5,959,435 | 6.09 | 6.65 | 5.13 | CHR16_5620349 | 17.5 | 47 |
|  | 18 | (*E*)-methyl hex-2-enoate | 12 | 9,262,853 | 11,969,757 | 29.29 | 35.49 | 8.02 | CHR12_10293679 | 25.9 | 259 |
|  | 19 | Ethyl acetate | 12 | 8,693,726 | 10,063,473 | 27.14 | 29.29 | 5.56 | CHR12_9262853 | 18.8 | 160 |
|  | 20 | (*E*)-ethyl but-2-enoate | 10 | 8,393,861 | 10,470,145 | 15.36 | 21.1 | 4.98 | CHR10_9422814 | 17 | 194 |
|  | 21 | Pentyl acetate | 18 | 2,190,801 | 3,906,155 | 4.91 | 7.22 | 5.07 | CHR18_2243935 | 17.3 | 153 |
| **Hydrocarbon** | 22 | Tetradecane | 2 | 11,363,813 | 14,717,925 | 47.83 | 54.29 | 5.14 | CHR02_13179708 | 17.5 | 100 |
| **Ketone** | 23 | 4,6-dimethyl-heptan-2-one | 14 | 5,819,037 | 10,973,137 | 11.07 | 19.48 | 5.69 | CHR14_6754203 | 19.2 | 235 |
|  | 24 | (*E*)-methyl hex-2-enoate | 2 | 20,300,502 | 21,206,376 | 64.07 | 65.22 | 6.19 | CHR02_20852996 | 20.7 | 48 |
|  | 25 | Octan-3-one | 14 | 18,823,692 | 23,119,473 | 29.94 | 38.05 | 5.73 | CHR14_23482094 | 19.3 | 203 |
| **Monoterpene** | 26 | Geraniol | 6 | 4,724,259 | 6,544,650 | 19.05 | 27.79 | 11.65 | CHR06_4994369 | 35.3 | 188 |
|  | 27 | Geraniol | 12 | 8,138,436 | 11,657,909 | 27.14 | 35.49 | 5.74 | CHR12_8540953 | 19.3 | 354 |
|  | 28 | Geraniol | 15 | 15,798,851 | 17,017,873 | 44.67 | 51.84 | 6.6 | CHR15_16157018 | 21.9 | 136 |
| | 29 | (*E*)-β-ocimene | 6 | 5,042,115 | 7,687,998 | 20.61 | 29.24 | 5.06 | CHR06_6544650 | 17.2 | 287 |
|  | 30 | (*E*)-methyl geranate | 12 | 10,063,405 | 15,681,332 | 29.29 | 41.56 | 6.49 | CHR12_11337672 | 21.6 | 336 |
|  | 31 | (*E*)-methyl geranate | 15 | 12,931,290 | 15,408,224 | 28.46 | 42.51 | 4.94 | CHR15_13509146 | 16.9 | 219 |
|  | 32 | Hotrienol | 2 | 5,077,251 | 6,341,438 | 27.28 | 33.3 | 5.58 | CHR02_5291224 | 18.9 | 126 |
|  | 33 | Hotrienol | 10 | 4,774,999 | 7,018,428 | 12.87 | 12.87 | 6.38 | CHR10_6908938 | 21.3 | 219 |
|  | 34 | Nerol | 10 | 3,870,491 | 5,568,232 | 12.85 | 12.87 | 6.76 | CHR10_3995724 | 22.4 | 206 |
| | 35 | γ-Isogeraniol | 15 | 9,619,429 | 13,331,325 | 16.86 | 33.87 | 5.39 | CHR15_11680187 | 18.3 | 258 |
| | 36 | β-Citronellol | 6 | 3,258,658 | 6,277,243 | 13.19 | 29.24 | 7 | CHR06_4994369 | 23 | 297 |
|  | 37 | (3*S*)-Linalool | 2 | 5,026,521 | 5,375,133 | 27.28 | 27.64 | 11 | CHR02_5198993 | 33.8 | 41 |
|  | 38 | (3*S*)-Linalool | 10 | 3,972,907 | 7,475,551 | 12.85 | 13.98 | 10.46 | CHR10_5431663 | 32.4 | 368 |
| **Norisoprenoid** | 39 | (*Z*)-Theaspirane | 1 | 3,653,671 | 4,103,100 | 15.75 | 17.18 | 25.07 | CHR01_4060900 | 60.9 | 36 |
|  | 40 | Dihydroactinidiolide | 17 | 13,721,527 | 24,127,437 | 48.78 | 56.64 | 11.11 | CHR17_24082626 | 34 | 327 |
|  | 41 | Cistus cyclohexanone | 11 | 316,912 | 783,312 | 1.47 | 2.88 | 5.46 | CHR11_338773 | 18.5 | 55 |
|  | 42 | Cistus cyclohexanone | 12 | 1,668,330 | 6,362,655 | 8.64 | 20.09 | 5.8 | CHR12_1668330 | 19.5 | 255 |
|  | 43 | Cistus cyclohexanone | 18 | 6,636,103 | 7,977,021 | 20.09 | 25.92 | 7.43 | CHR18_6525848 | 24.3 | 153 |
| Sesquiterpene | 44 | (Z,E)-α-Farnesene | 9 | 7,083,270 | 7,644,375 | 35.29 | 35.29 | 7.25 | CHR09_7215558 | 23.8 | 30 |
| | 45 | (Z,E)-α-Farnesene | 14 | 28,849,280 | 31,295,985 | 58.79 | 63.8 | 5.12 | CHR14_30214461 | 17.4 | 295 |
| | 46 | (Z,E)-α-Farnesene | 1 | 5,204,981 | 6,064,966 | 21.82 | 22.89 | 14.59 | CHR01_5394295 | 42.1 | 83 |
|  | 47 | Dihydroedulan | 18 | 2,147,296 | 4,008,254 | 4.91 | 6.85 | 11.01 | CHR18_2688697 | 33.8 | 173 |
|  | 48 | Germacrene D | 18 | 1,441,513 | 3,020,876 | 3.68 | 4.91 | 8.94 | CHR18_2688697 | 28.4 | 125 |
| | 49 | δ-Cadinene | 2 | 6,997,965 | 9,134,829 | 37.34 | 41.37 | 5.55 | CHR02_7523961 | 65.5 | 125 |
| Monoterpene Glycoside | 50 | PH1 | 13 | 20,312,618 | 22,962,230 | 47.37 | 51.7 | 23.93 | CHR13_22872934 | 61.6 | 119 |
|  | 51 | PH2 | 13 | 20,313,457 | 22,962,230 | 47.37 | 51.7 | 29.08 | CHR13_21136279 | 68.8 | 137 |
|  | 52 | PH3 | 13 | 19,225,568 | 23,236,187 | 44.51 | 52.77 | 17.54 | CHR13_19871765 | 50.5 | 172 |
|  | 53 | PH4 | 13 | 16,798,759 | 22,962,230 | 43.8 | 51.7 | 14.68 | CHR13_19603418 | 44.4 | 254 |
|  | 54 | PH5 | 2 | 5,077,251 | 5,566,371 | 27.28 | 27.64 | 8.01 | CHR02_5198993 | 27.4 | 46 |
|  | 55 | PH5 | 10 | 3,995,724 | 5,748,943 | 12.86 | 12.87 | 19.15 | CHR10_5486459 | 53.6 | 162 |
|  | 56 | PH6 | 2 | 1,379,492 | 6,341,438 | 8.16 | 33.3 | 6.87 | CHR02_2008551 | 24.1 | 544 |
|  | 57 | PH6 | 10 | 3,903,774 | 8,393,861 | 12.85 | 15.36 | 5.75 | CHR10_5422073 | 20.6 | 467 |
|  | 58 | PH6 | 15 | 14,702,159 | 16,690,679 | 42.51 | 48.99 | 7.01 | CHR15_16157018 | 24.5 | 205 |
|  | 59 | PH8 | 5 | 22,463,101 | 24,772,382 | 53.93 | 57.42 | 5.54 | CHR05_24327860 | 19.9 | 124 |

Most sesquiterpenes and all three norisoprenoids were associated with significant QTLs. (*Z*)-theaspirane and dihydroactinidiolide mapped to single loci on Chr 1 and 17, respectively, whereas cistus cyclohexanone was associated with three QTLs on Chr 11, 12, and 18. Among sesquiterpenes, significant QTLs were detected for (*E*)-β-farnesene on Chr 1, 9, and 14, while cyclic sesquiterpenes mapped to Chr 2 and 19. Overlapping QTLs on Chr 18 were identified for germacrene D and dihydroedulan, and δ-cadinene was associated with a single major QTL on Chr 2.

Of the 12 monoterpenes detected, eight were associated with significant QTLs on Chr 2, 6, 10, 12, and 15. Geraniol and its derivatives (E-methyl geranate and γ-isogeraniol) colocalized on Chr 15, with an additional shared QTL on Chr 12. Geraniol, (*E*)-β-ocimene, and β-citronellol mapped to the same region on Chr 6. QTLs associated with the abundance of (3*S*)-linalool, hotrienol, and nerol colocalized on Chr 10, and (3S)-linalool and hotrienol also shared a QTL on Chr 2. Both (3*S*)-linalool QTLs were major, explaining 33.8% and 32.4% of the variation.

Seven of the 14 monoterpene glycosides were associated with significant QTLs. PH1-PH4 colocalized on Chr 13, despite the absence of QTLs for free monoterpenes in this region. In contrast, PH5 mapped to two major QTLs overlapping those identified for (3*S*)-linalool, jointly explaining 81% of phenotypic variation, with the Chr 10 locus alone accounting for 53.6%. Given the major QTL effects and its central role in Riesling aroma, subsequent analyses focused on the genetic basis of (3*S*)-linalool production.

### Confirmation of QTL effects on linalool and linalool glycoside abundance

Two major QTLs were identified for the accumulation of free (3*S*)-linalool and the highly correlated glycoside PH5 on Chr 2 and 10 (**Fig. 6A**). The Chr 2 QTL spanned ∼0.34 Mb for both traits (0.35 cM), whereas the Chr 10 QTLs were broader, spanning 3.5 Mb for (3S)-linalool and 1.8 Mb for PH5 (1.13 cM). To assess allelic effects, we examined the relationship between compound abundance and genotype at peak markers within each QTL. On Chr 2, both traits shared the same peak marker (CHR02_5198993). On Chr 10, peak markers differed slightly (CHR10_5431663 for (3*S*)-linalool and CHR10_5486459 for PH5) within a broad interval encompassing >200 markers above the genome-wide significance threshold.

**Figure 6.**
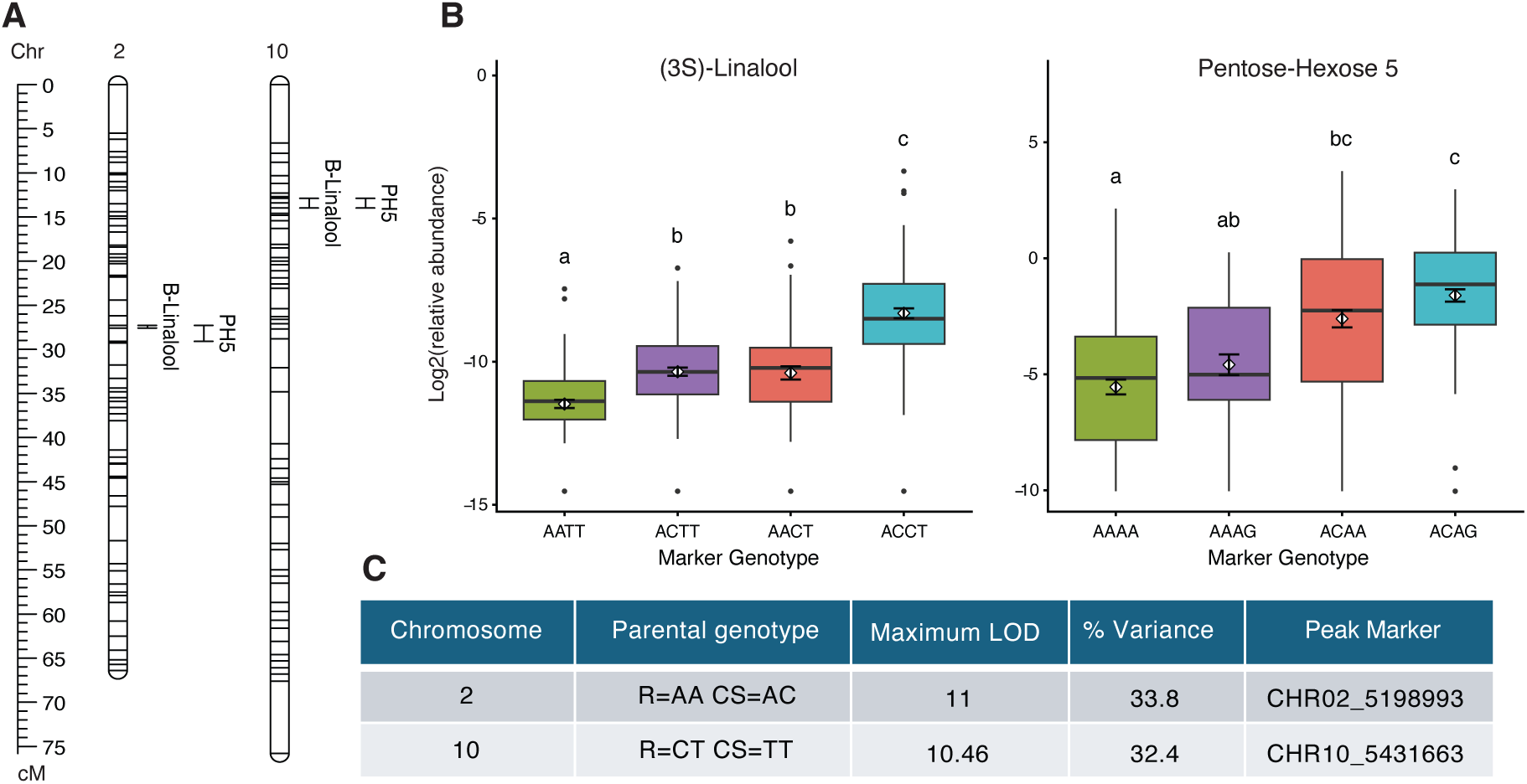
Marker effects of (3*S*)-linalool QTLs in the RxCS population. (**A**) Genetic maps of chromosomes 2 and 10 showing the positions of QTLs for (3*S*)-linalool and pentose-hexose glycoside 5 (PH5), with 95% LOD confidence intervals. (**B**) Distribution of (3*S*)-linalool (left) and PH5 (right) relative abundance across the RxCS population, grouped by genotype at peak markers within each QTL interval. Relative abundances for (3*S*)-linalool and PH5 are in octan-2-ol and decyl-β-D-glucopyranoside equivalents, respectively. Boxplots show median and interquartile range; points represent individual genotypes. Different letters indicate significant differences among genotype classes (Tukey’s HSD, *P* < 0.05). (**C**) Summary of QTL statistics for (3*S*)-linalool, including chromosome location, parental genotype configuration, maximum LOD score, percentage of explained variance, and peak marker.

Linear models were used to estimate the proportion of variance explained by the markers. Modeling revealed strong additive effects and modest but significant interactions between markers. For (3*S*)-linalool, the additive model explained 38.4% of phenotypic variance, increasing to 44.4% when interactions were included. For PH5, marker effects were stronger, with the additive model explaining 61.8% of the variance. In both cases, the Chr 10 locus contributed more to phenotypic variation than the Chr 2 locus. For (3*S*)-linalool, CHR10_5431663 and CHR02_5198993 showed strong effects (F = 87.1 and 69.9, respectively; *P* < 10⁻¹⁵). For PH5, CHR10_5431663 was the primary predictor (F = 322.0; *P* = 8.76 x 10⁻⁵⁰), followed by CHR02_5198993 (F = 87.1; *P* = 2.08 x 10⁻¹⁸). Given that the aglycone of PH5 was not experimentally confirmed, subsequent analyses focused on loci associated with free (3*S*)-linalool.

Genotype effects were consistent with additive contributions from both loci. Individuals heterozygous at both loci (AC/CT at CHR02_5198993 and CHR10_5431663) exhibited the highest (3*S*)-linalool levels, significantly exceeding all other genotype combinations. Heterozygosity at a single locus (AC/TT or AA/CT) resulted in intermediate levels (**Fig. 6B**). Riesling is homozygous (AA) at CHR02_5198993, whereas Cabernet Sauvignon is homozygous (TT) at CHR10_5431663, suggesting that the C allele at these loci is derived from RHap2 (**Fig. 6C**). A subset of homozygous and heterozygous genotypes was selected for transcriptome and metabolite profiling in 2021.

### Integration of transcriptome and metabolite data to identify candidate genes for linalool accumulation

To aid in the identification of candidate genes associated with linalool accumulation, we sequenced berry transcriptomes of both parents and eight progeny in fall 2021 at two time points: 14 weeks post-anthesis (preharvest) and at harvest (15-17 weeks post-anthesis). Progeny were selected based on marker genotype and fruit pigmentation. At preharvest stage, 18,789 genes were differentially expressed between Riesling and Cabernet Sauvignon (|log2FC| > 1, adjusted *P* < 0.01), increasing to 20,490 at harvest (**Supplementary Table S4**).

Berry samples collected at the same time points were analyzed for volatile and glycosylated compounds using HS-SPME-GC-MS and UHPLC-QTOF/MS. A total of 77 volatile compounds were detected, of which 65 differed significantly among genotypes (**Supplementary Table S5A, Figure 7**). 13 monoterpene glycosides were detected (**Supplementary Table S5B, Figure 7**). All compounds were detected at both developmental stages. Hierarchical clustering based on monoterpenoid and monoterpene glycoside abundance separated genotypes into three groups. Genotypes heterozygous (CT) at the CHR10_5431663 marker formed distinct clusters from homozygous individuals, although the individuals 95 and 106 grouped closer to TT genotypes. Nearly all monoterpenoid glycoside abundances were strongly correlated between years (r^2^ = 0.83-0.97, adjusted *P* < 0.003, **Supplementary Table S6A**). Fifty-five volatile compounds were identified in both 2020 and 2021. With an r^2^ value of 0.92 and adjusted *P* value of 0.0002, (3*S*)-linalool was also highly correlated between seasons, indicating consistent genotype-dependent abundance across years (**Supplementary Table S6B**). Over half of the other volatile compounds were moderately to strongly correlated between years (r^2^ = 0.42-0.76) but many correlations did not remain significant after correcting for multiple testing (**Supplementary Table S6B**). A total of 31 compounds differed significantly between preharvest and harvest stages, with most increasing in abundance at harvest (**Supplementary Fig. S3**, **Supplementary Table S7**). Only five compounds were more abundant at preharvest, including the norisoprenoid (*Z*)-theaspirane and four lipoxygenase-derived compounds (e.g., (*Z*)-3-hexen-1-ol).

**Figure 7.**
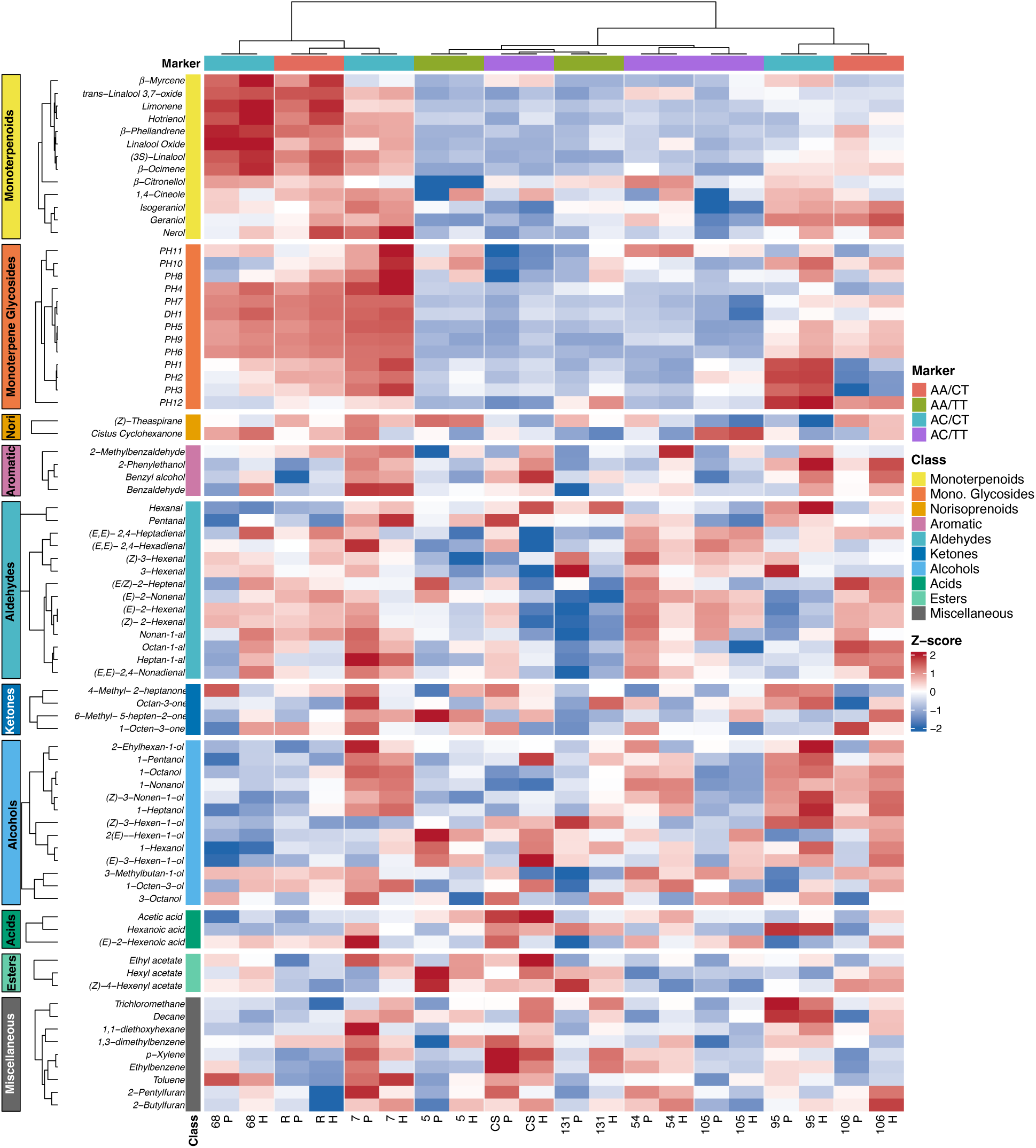
Z-score heatmap of volatile compounds and monoterpene glycosides in RxCS berry samples. Heatmap showing standardized (Z-score) relative abundances of volatile compounds and monoterpene glycosides across genotypes (columns). Rows (compounds) are grouped and hierarchically clustered within compound classes, and columns are clustered based on monoterpenoid and monoterpene glycoside profiles. Genotypes are annotated by marker genotype at the Chr 2 and Chr 10 (3*S*)-linalool QTL peak markers. Color scale indicates relative abundance (red, higher; blue, lower).

To integrate metabolite and transcriptome data, volatile and glycosylated monoterpenoid abundances were combined with gene expression data within the Chr 10 (3*S*)-linalool QTL interval across all four haplotypes using multiple factor analysis (MFA). The first two dimensions explained 32.9% and 17.4% of the total variance, respectively (**Fig. 8A)**. Individuals formed three clusters corresponding to marker genotypes at the Chr 2 and Chr 10 QTLs, with genotypes homozygous at CHR10_5431663 grouping together. The variable projection (**Fig. 8B**) showed strong alignment of monoterpenoids and monoterpene glycosides with multiple genes from RHap2, indicating coordinated variation between gene expression and metabolite accumulation. Among these, *VITVvi_vRiesFPS24_v1.1.hap2.chr10.ver1.1.g472680*, annotated as a (3*S*)-linalool/-nerolidol synthase, showed the strongest association with monoterpenoid abundance. In contrast, most genes from Cabernet Sauvignon haplotypes and RHap1 were projected in the opposite direction, suggesting haplotype-specific regulation.

**Figure 8.**
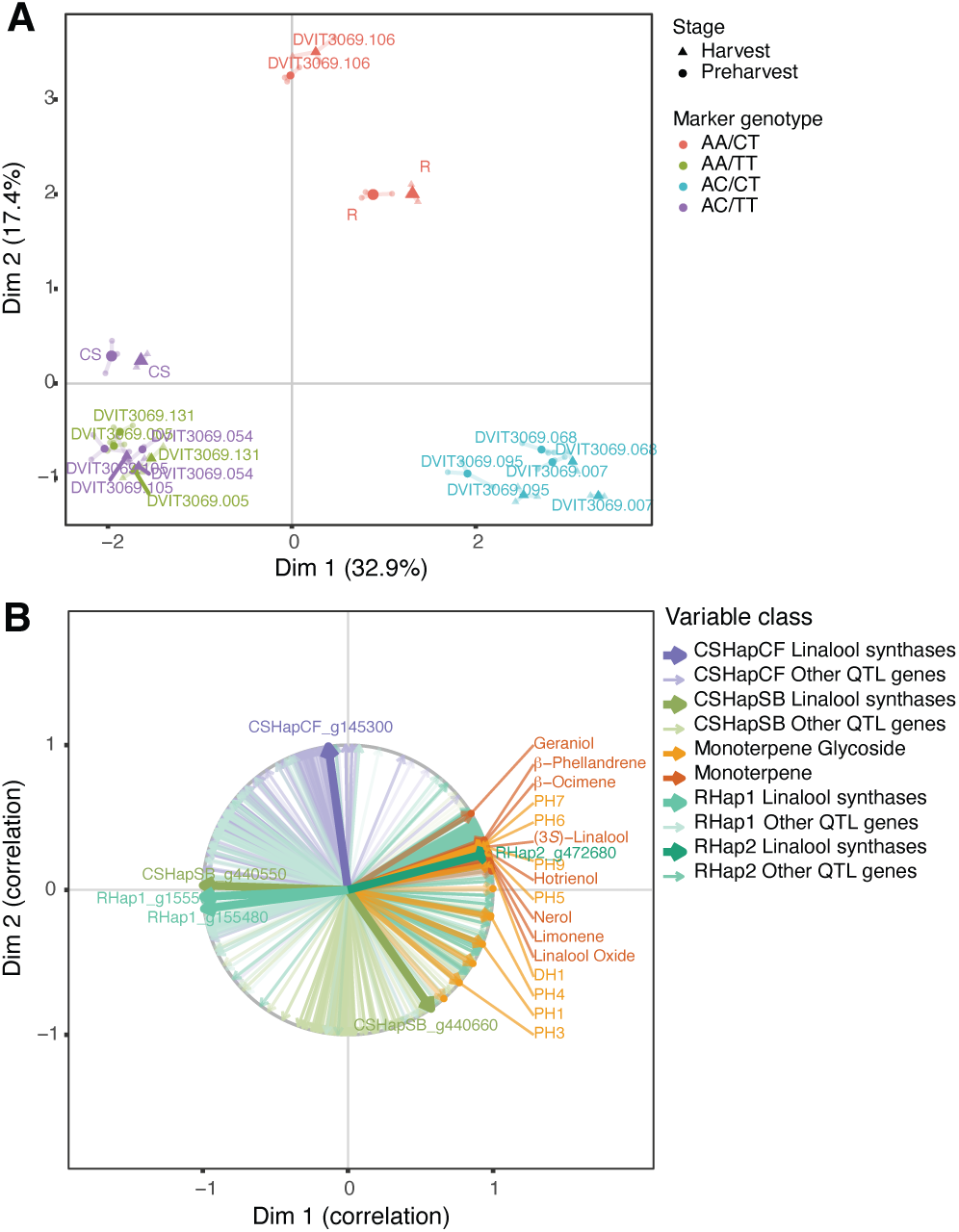
Multiple factor analysis (MFA) integrating monoterpenoids, monoterpene glycosides, and haplotype-specific gene expression within the linalool QTL on chromosome 10. (**A**) MFA individuals plot showing the distribution of RxCS progeny. Points represent individual samples, colored by marker genotype at Chr 2 and Chr 10 QTLs and shaped by developmental stage (preharvest or harvest). The first two dimensions explain 32.9% and 17.4% of total variance, respectively. (**B**) MFA variables plot showing correlations between metabolites and gene expression variables within the Chr 10 QTL interval. Only the top contributing monoterpenoids and monoterpene glycosides are labeled for clarity. Genes encoding (3*S*)-linalool/(*E*)-nerolidol synthases are highlighted in bold. Variable classes are color-coded by haplotype and functional category.

We further assessed gene-metabolite relationships using Spearman correlation with (3*S*)-linalool content. Given previous reports linking MEP pathway genes to linalool accumulation in Muscat cultivars (Wang *et al*., 2021), we initially examined this pathway. However, *VviDXS1* expression was not significantly correlated with linalool levels. Only two genes showed weak positive correlations: CDP-ME kinase (*CMK*; *VITVvi_vRiesFPS24_v1.1.hap2.chr06.ver1.0.g409230*) and geranyl-diphosphate synthase (*GPPS*; *VITVvi_vRiesFPS24_v1.1.hap1.chr15.ver1.0.g235820*), at harvest and preharvest, respectively (**Supplementary Table S8**).

### Identification of a candidate TPS gene within the Chr 10 (3S)-linalool QTL

The number of annotated genes within the Chr 10 QTL interval ranged from 299 to 397 across haplotypes (**Fig. 9A**). Each haplotype contained a cluster of 10-12 terpene synthase genes, annotated as (3S)-linalool/nerolidol synthases. After filtering out lowly expressed genes (TPM < 2 in at least 12 samples), 109-124 genes remained expressed within the interval across haplotypes. Spearman correlation analysis revealed marked differences among haplotypes in the number of genes associated with (3*S*)-linalool content (**Fig. 9B**). Ten or fewer genes from RHap1 and both Cabernet Sauvignon haplotypes (Cochetel *et al*., 2025) showed significant correlations, whereas 66 genes from RHap2 were significantly correlated, including 36 with strong associations (ρ > 0.7; adjusted *P* < 4.2 x 10^-9^; **Fig. 9C; Supplementary Table S9**). Genes with higher expression variance across samples tended to show stronger correlations, suggesting that transcriptional variability contributes to metabolite variation within the population.

**Figure 9.**
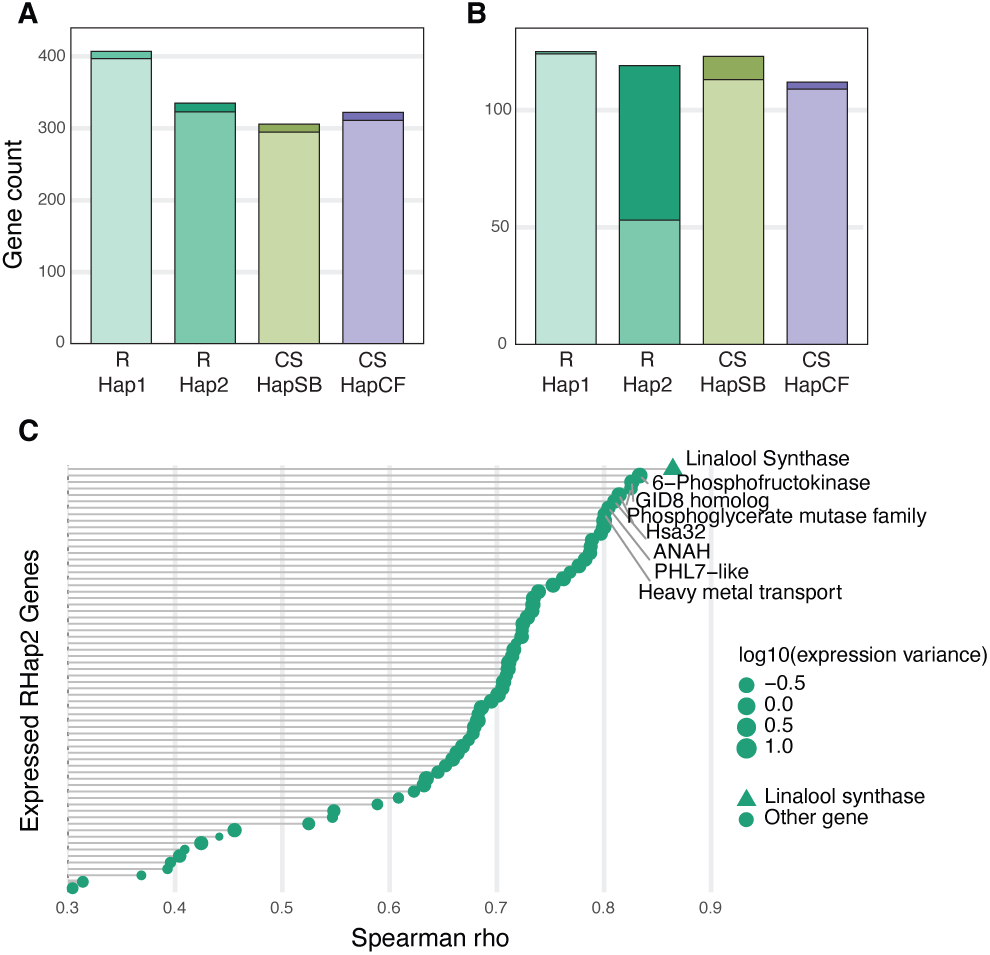
Identification of candidate genes associated with linalool accumulation within the chromosome 10 QTL interval. (**A**) Number of genes located within the Chr 10 QTL interval for each parental haplotype (RHap1, RHap2, CSHapSB, CSHapCF). (**B**) Number of genes expressed in berry tissue (TPM ≥ 2 in at least 12 samples) within each haplotype-specific interval. Darker segments indicate genes significantly correlated with linalool abundance (Spearman; adjusted *P* < 0.05). (**C**) Spearman correlation between gene expression and (3*S*)-linalool abundance for expressed RHap2 genes (TPM ≥ 2 in at least 12 samples). Each point represents a gene ranked by correlation coefficient (ρ). Point size reflects log_10_-transformed expression variance across samples. Genes with ρ > 0.3 and adjusted *P* < 0.05 are shown; the top eight genes are labeled with functional annotations.

The gene with the highest correlation across time points was *VITVvi_vRiesFPS24_v1.1.hap2.chr10.ver1.1.g472680* (ρ = 0.86; adjusted *P* = 2.3 × 10⁻¹⁶), annotated as a (3*S*)-linalool/nerolidol synthase. Both Spearman correlation and MFA independently identified this gene as the strongest candidate within the Chr 10 QTL. Although several neighboring genes showed similarly high correlation coefficients, they lack functional annotation related to terpene biosynthesis and are likely co-inherited within the same haplotype block. In contrast, none of the 20-25 genes in the Chr 2 QTL of each haplotype was significantly associated with (3*S*)-linalool (**Supplementary Table S10**).

Alignment of the VITVvi_vRiesFPS24_v1.1.hap2.chr10.ver1.1.g472680 protein sequence against the monoterpene synthases annotated in *V. v. vinifera* PN40024 line showed 98.2% sequence identity with VIT_00s0385g00020 (VviTPS54/VviPNLinNer1), a functionally characterized (3*S*)-linalool/(*E*)-nerolidol synthase belonging to the TPS-g subfamily (VviTPS54 in Martin *et al*., 2010) (**Supplementary Table S11**). Multiple sequence alignment with Cabernet Sauvignon homologs and VIT_00s0385g00020 revealed high conservation at the amino acid level, with no major structural differences. However, the RHap1 homolog (VITVvi_vRiesFPS24_v1.1.hap1.chr10.ver1.1.g155480) appeared truncated at the N-terminus.

Analysis of N-terminal sequences across all four homologs did not identify a plastid-targeting peptide (**Supplementary Table S12**), consistent with previous observations that targeting signals are poorly conserved in grape TPS-g proteins despite their predicted plastid localization (Martin *et al*., 2010). All sequences retained the conserved DDxxD and NSE/DTE motifs characteristic of terpene synthases (**Supplementary Figure S4**) (Degenhardt *et al*., 2009).

Together, these results indicate that functional differences in (3*S*)-linalool accumulation are unlikely to be driven by major changes in protein sequence. Instead, the strong association between (3*S*)-linalool levels and expression of *VITVvi_vRiesFPS24_v1.1.hap2.chr10.ver1.1.g472680* suggests that transcriptional regulation is a key driver of the phenotype. To explore potential regulatory mechanisms, we analyzed transcription factor-binding sites within 3 kb up- and downstream of the gene across haplotypes (**Supplementary Table S13**). Although differences in predicted binding site composition were observed for the RHap2 allele, the distribution of binding sites for known regulators of grape monoterpenoid biosynthesis such as bHLH and MYB family members was similar for the majority of haplotypes (Li *et al*., 2024; Sun *et al*., 2025). However, correlation analysis identified several transcription factor genes differentially expressed between Riesling and Cabernet Sauvignon that were strongly associated with both (3*S*)-linalool content and expression of *VITVvi_vRiesFPS24_v1.1.hap2.chr10.ver1.1.g472680* (**Table 4**).

**Table 4:** Top 10 transcription factor genes exhibiting significant Spearman correlation (ρ) with (3*S*)-linalool content. All genes are significantly differentially expressed between Riesling and Cabernet Sauvignon.

| Gene ID | Function | $\rho$ | Adjusted *P* |
| --- | --- | --- | --- |
| VITVvi_vRiesFPS24_v1.1.hap1.chr10.ver1.0.g159690 | trihelix transcription factor ASIL2-like | 0.77 | 2.69 x10<sup>-3</sup> |
| VITVvi_vRiesFPS24_v1.1.hap1.chr10.ver1.0.g162350 | S-adenosyl-L-methionine-dependent methyltransferases superfamily | 0.79 | 1.20 x10<sup>-4</sup> |
| VITVvi_vRiesFPS24_v1.1.hap1.chr19.ver1.0.g303420 | Probable WRKY transcription factor 31 | 0.79 | 4.30 x10<sup>-4</sup> |
| VITVvi_vRiesFPS24_v1.1.hap2.chr08.ver1.0.g434550 | Transducin WD40 repeat-like superfamily | 0.85 | 3.50 x10<sup>-4</sup> |
| VITVvi_vRiesFPS24_v1.1.hap2.chr08.ver1.0.g434460 | MYB SANT-like DNA-binding domain | 0.84 | 1.00 x10<sup>-4</sup> |
| VITVvi_vRiesFPS24_v1.1.hap2.chr10.ver1.0.g471280 | Heavy metal transport detoxification superfamily | 0.82 | 3.90 x10<sup>-4</sup> |
| VITVvi_vRiesFPS24_v1.1.hap2.chr10.ver1.0.g472350 | transcription factor ILR3-like | 0.8 | 1.10 x10<sup>-3</sup> |
| VITVvi_vRiesFPS24_v1.1.hap2.chr10.ver1.0.g471950 | MYB family transcription factor PHL7-like | 0.79 | 1.76 x10<sup>-3</sup> |
| VITVvi_vRiesFPS24_v1.1.hap2.chr10.ver1.0.g474280 | Transcription factor | 0.76 | 3.30 x10<sup>-3</sup> |
| VITVvi_vRiesFPS24_v1.1.hap2.chr19.ver1.0.g623080 | probable RNA polymerase II transcription factor B subunit 1-1 | 0.77 | 4.00 x10<sup>-4</sup> |

## Discussion

The RxCS population segregated for a wide range of volatile compounds, including key grape monoterpenoids, sesquiterpenes, norisoprenoids, and phenylpropanoid derivatives. QTL analysis identified 49 significant loci associated with 40 volatile compounds, with an additional 10 loci linked to seven monoterpene glycosides. As expected, Riesling accumulated substantially higher monoterpenoid levels than Cabernet Sauvignon (Kalua and Boss, 2010), and several compounds exhibited transgressive segregation, consistent with multigenic control of volatile biosynthesis in grape (Doligez *et al*., 2006; Bosman *et al*., 2023). QTLs for structurally or biosynthetically related compounds frequently colocalized, suggesting shared genetic control at multiple steps within biosynthetic pathways.

Much of the genetic variation in monoterpene accumulation in grape has been attributed to a gain-of-function mutation in *VviDXS1* on Chr 5, which increases precursor supply through the MEP pathway (Battilana *et al*., 2011). However, monoterpene biosynthesis is clearly multigenic, with multiple QTLs identified beyond this locus across different populations (Doligez *et al*., 2006; Battilana *et al*., 2009; Emanuelli *et al*., 2010; Dalla Costa *et al*., 2018; Bosman *et al*., 2023). Notably, 75% of aromatic non-Muscat cultivars lack functional *VviDXS1* mutations, indicating that other loci make substantial contributions to monoterpenoid accumulation (Emanuelli *et al*., 2010). In the RxCS population, no QTL on Chr 5 was associated with monoterpenoid content, and *VviDXS1* expression was not significantly correlated with terpene levels. This is consistent with prior work showing that introduction of the Muscat *VviDXS1* allele increases some monoterpenes without inducing linalool synthase expression or linalool accumulation (Dalla Costa *et al*., 2018) and with genetic evidence that linalool content can be inherited independently of total monoterpene content (Duchêne *et al*., 2009a). Together, these findings support a two-step model in which precursor supply and monoterpene composition are controlled by distinct loci and confirm that Riesling’s intermediate monoterpenoid accumulation is not explained by a *VviDXS1* gain-of-function allele.

The two major (3*S*)-linalool QTLs on Chr 2 and Chr 10 overlap loci consistently reported across diverse grapevine populations (Doligez *et al*., 2006; Battilana *et al*., 2009; Duchêne *et al*., 2009a; Koyama *et al*., 2022), suggesting they represent conserved genetic determinants of linalool accumulation. Each QTL explained more than 30% of phenotypic variation. Linear modeling of corresponding marker genotypes further showed that the markers accounted for 38.4% of variance in an additive model, increasing to 44.4% when marker interactions were included. Despite their repeated identification, candidate genes within these intervals had not previously been proposed, likely due to the highly duplicated and structurally variable nature of TPS genes and the limitations of haploid reference genomes in resolving heterozygous loci.

Terpene synthases comprise a large and structurally complex gene family in grape, with 80 to over 200 members per cultivar (Martin *et al*., 2010; Bosman and Lashbrooke, 2023; Bosman *et al*., 2023). High heterozygosity and tandem duplication have historically hindered placement of individual TPS genes, leaving key linalool synthases *VviTPS54* and *VviTPS56* unplaced in earlier reference genomes (Martin *et al*., 2010; Smit *et al*., 2020). Using phased diploid assemblies, a *TPS-g* gene cluster including *VviTPS54* and *VviTPS56* was resolved on Chr 10 (Smit *et al*., 2020), corresponding precisely to the Chr 10 QTL identified here. Functional assays have shown that *VviTPS56* alone is sufficient to produce (3*S*)-linalool, with *VviDXS1* coexpression increasing yield but not being required (Wang *et al*., 2021), supporting a model in which downstream terpene synthase activity, rather than *VviDXS1*-driven precursor supply, is the primary determinant of linalool accumulation in non-Muscat aromatic cultivars.

To identify candidate genes within the Chr 10 interval, we examined correlations between linalool and haplotype-resolved gene expression across the QTL region. The number of genes significantly correlated with (3*S*)-linalool content differed dramatically between haplotypes: 66 RHap2 genes showed significant correlation compared to ten or fewer in RHap1 and both Cabernet Sauvignon haplotypes. The gene with the highest correlation coefficient across both timepoints was the RHap2 homolog of *VviTPS54/VviPNLinNer1*, a functionally characterized (3*S*)-linalool/(*E*)-nerolidol synthase (Martin *et al*., 2010), a result corroborated by MFA. In Riesling, *VviTPS54* expression correlates with linalool content while *VviTPS56* declines toward the end of ripening (Yue *et al*., 2020), and both genes are expressed during later stages of berry development in parallel with linalool accumulation in aromatic cultivars (Matarese *et al*., 2013; Wen *et al*., 2015), further supporting *VviTPS54* as the primary driver of linalool accumulation at harvest.

Protein sequence alignment across all four haplotypes revealed high conservation at key functional sites of *VviTPS54* (**Supplementary Figure 5**), including the DDxxD and NSE/DTE metal-binding motifs, with no major substitutions predicting altered enzymatic activity (Kampranis *et al*., 2007; Drew *et al*., 2016). The RHap1 homolog appeared truncated at the N-terminal domain, likely rendering it non-functional. Together these observations suggest that sequence divergence does not explain linalool accumulation differences between Riesling and Cabernet Sauvignon. Instead, the strong haplotype-specific correlation between *VviTPS54* transcript abundance and linalool content points to regulatory variation as the primary mechanism. Consistent with this, exogenous salicylic acid increased *VviTPS54* expression and (3S)-linalool content in Muscat Hamburg (Yue *et al*., 2023), overexpression of *VvbZIP61* in *V. amurensis* calli increased monoterpenoid production, and *VvbZIP61* was differentially expressed in aromatic versus non-aromatic cultivars (Zhang *et al*., 2023), suggesting that trans-acting regulators of TPS expression vary between cultivars.

The molecular basis for elevated *VviTPS54* expression in RHap2 remains unresolved. Transcription factor-binding site analysis in the 3 kb flanking regions of all four homologs identified differences proximate to RHap2, but none with established roles in monoterpenoid regulation (Li *et al*., 2024; Sun *et al*., 2025). Several transcription factor genes differentially expressed between Riesling and Cabernet Sauvignon correlated with both (3*S*)-linalool content and *VviTPS54* expression, suggesting possible trans-regulatory control. Transcription factors located on Chr 2 belonging to the bHLH family have been shown to correlate with (3S)-linalool accumulation; however, these lie outside the QTL interval (Sun *et al*., 2025). Epigenetic mechanisms such as differential DNA methylation could also contribute to haplotype-specific expression and cultivar- or clone-specific variation in monoterpenoid accumulation (Cui *et al*., 2026), and copy number variation within the Chr 10 TPS cluster may further contribute to differences in total linalool synthase activity (Bosman *et al*., 2023).

Although no candidate gene was identified within the Chr 2 interval, this locus has been independently detected across multiple populations (Sevini *et al*., 2004; Doligez *et al*., 2006; Battilana *et al*., 2009; Duchêne *et al*., 2009a; Koyama *et al*., 2022) and made a significant additive contribution alongside Chr 10. One possibility is that it acts through a regulatory mechanism operating at an earlier developmental stage than those sampled here; additional time points spanning early berry development could help clarify its role. We observed a slight bias toward higher (3*S*)-linalool in unpigmented genotypes, suggesting a possible link to anthocyanin biosynthesis. However, *VviMYBA1* and *VviMYBA2* also map outside the Chr 2 QTL interval, and the same locus was identified in selfed progeny of Muscat Ottonel (Duchêne *et al*., 2009a), which are all unpigmented, ruling out anthocyanin status as the source of variation at this locus. The bias toward unpigmented genotypes may instead reflect independent upregulation of linalool synthases: in Tempranillo blanco, *VviTPS54*, *55*, *58*, *60*, and *64* are upregulated relative to the pigmented Tempranillo tinto, with *VviTPS55* and *VviTPS60* also responding to UV-B during ripening (Carbonell-Bejerano *et al*., 2014; Rodríguez-Lorenzo *et al*., 2023). Whether anthocyanin status directly influences *VviTPS54* expression in the RxCS population could not be resolved with our RNA-Seq sample size and warrants further investigation.

The distribution of monoterpenoid content beyond (3*S*)-linalool reflects a more complex genetic architecture involving multiple loci and biosynthetic steps. Hotrienol and nerol oxide, which arise through enzyme-mediated oxidation and rearrangement of linalool (Williams *et al*., 1980; Luan *et al*., 2006), co-mapped with the Chr 2 and Chr 10 linalool QTLs, consistent with their metabolic relationship. Geraniol and its derivatives mapped to distinct loci on Chr 6, 12, and 15, indicating separate genetic control of different monoterpene end products. The strongly correlated glycoside PH5 shared both linalool QTLs, suggesting coordinated regulation of biosynthesis and glycosylation, while the Chr 13 glycoside QTL shared by PH1-4, lacking a corresponding free monoterpenoid QTL, may reflect variation in glycosyltransferase activity and should be addressed in future work.

Volatile QTL mapping in the full population was conducted using a single season of phenotypic data. Because volatile abundance in grape berries can be strongly influenced by environmental and vintage effects (Caven-Quantrill and Buglass, 2008; Lu *et al*., 2022), multi-year phenotyping would help evaluate QTL stability (Quero-García *et al*., 2021). However, linalool abundance measured in a second year in the transcriptome subset was strongly correlated with 2020 data, the detected QTLs overlap loci reported in other populations and environments, and the convergence of metabolite, transcriptomic, and comparative QTL evidence supports the Chr 10 locus as a recurrent genetic determinant of linalool variation rather than a season-specific artifact.

This study links (3*S*)-linalool accumulation in an intraspecific wine grape population to a terpene synthase cluster on Chr 10 and identifies *VviTPS54* as a strong candidate gene driven by regulatory rather than coding sequence variation. The phased Riesling genome assembly was critical for resolving this highly duplicated and heterozygous locus and demonstrates the value of haplotype-resolved references for dissecting complex quantitative traits in grape. Together, the integration of untargeted volatile metabolomics, high-density QTL mapping, and haplotype-resolved transcriptomics provides a framework for marker-assisted selection of aroma traits and for understanding monoterpenoid diversity across wine grape cultivars.

## Acknowledgments

We thank the UC Davis Genome Center DNA Technologies Core Facility for their assistance with sequencing; Dr Andrea Minio for his bioinformatics support; Gregory Stevens and Olivia Haley for their help with viticultural operations and sampling; and Guillermo Garcia Zamora for vineyard management.

## Author contribution

DC and JL: conceptualization; DC and SE: supervision; JL and DC: writing; DC: funding acquisition; JL: experimentation; JL: data analysis and visualization; JL, MD-L: sampling; JL, MM, NC, LD-G: bioinformatic analysis; LL, JL: chemical analysis; RF-B: library preparation.

## Data Availability

All raw RNA-seq data are available on NCBI SRA under BioProject PRJNA1452220. The phased diploid genome assembly of Riesling and its structural annotation are deposited on NCBI under BioProject numbers PRJNA1453061 (Haplotype 1 and unplaced sequences) and PRJNA1453060 (Haplotype 2). Mass spectrometry data are available on Zenodo (DOI: 10.5281/zenodo.19747775).

